# Systemic dissemination, reproductive tissue tropism and gestational stage-dependent placental responses during Mayaro virus infection in a murine model

**DOI:** 10.64898/2026.01.04.693264

**Authors:** A.P. Arévalo, P. Perbolianachis, J.L. Pórfido, M. Pereira-Gómez, G. Greif, J. Hurtado, A. Fajardo, B. Varela, J.M. Verdes, G. Moratorio, M Crispo

## Abstract

Mayaro virus (MAYV) is an emerging mosquito-borne alphavirus associated with acute febrile illness and persistent arthralgia, with increasing reports in Latin America and potential for geographic expansion. However, its dissemination dynamics, tissue tropism and transmission routes remain incompletely defined. Here, we developed a murine model to characterize systemic viral dissemination, tissue tropism and the impact of gestational stage on maternal–fetal outcomes. A nanoluciferase-expressing MAYV reporter enabled non-invasive *in vivo* imaging, revealing rapid and widespread systemic dissemination under controlled conditions. Complementary infection with wild-type (WT) virus confirmed broad tropism across lymphoid, musculoskeletal and reproductive tissues. At 24 hours post-infection (hpi), viral distribution was relatively homogeneous across tissues, whereas at 48 hpi, tissue-specific differences emerged, with increased viral loads in selected organs, including the spleen and male reproductive tissues. Transient sex-dependent differences were observed at 24 hpi but were not sustained at later time points. Hematological and biochemical analyses revealed early systemic alterations consistent with changes in leukocyte distribution during acute infection. Infectious viral particles were detected in reproductive tissues of both sexes, including in sperm, supporting the biological plausibility of non-vector transmission, although detection in exposed animals was limited. Gestational stage influenced infection outcomes: early gestation (infected at 7.5–8.5 dpc; analyzed at 9.5–10.5 dpc) was associated with detection of infectious virus in fetal tissues, whereas mid-gestation (infected at 13.5–14.5 dpc; analyzed at 15.5–16.5 dpc) showed no detectable infectious virus in fetuses despite evidence of viral antigen persistence and sustained infectious viral presence in maternal and placental compartments. Together, these findings provide a preclinical framework for investigating MAYV pathogenesis and underscore the role of tissue tropism and gestational context in shaping infection dynamics and maternal–fetal involvement.

**Author summary:** Mayaro virus (MAYV) is an emerging mosquito-borne virus that causes fever and long-lasting joint pain, with increasing reports in Latin America and potential for wider geographic spread. Despite its growing relevance, key aspects of MAYV pathogenesis remain to be fully defined, including the mechanisms underlying systemic dissemination and tissue tropism, as well as its potential to affect pregnancy or be transmitted through non-vector routes. Here, we used a mouse model with a luminescent reporter virus to track MAYV dissemination *in vivo*, revealing rapid and widespread distribution across multiple tissues. In studies using wild-type virus, infectious virus was detected not only in muscle and immune-related organs but also in reproductive tissues of both females and males. Importantly, gestational stage influenced viral distribution. When infection occurred early in gestation, infectious viral particles were detected in fetal tissues. In contrast, at later stages of pregnancy, infectious virus was no longer detectable in fetuses, despite evidence of viral antigen persistence and sustained infectious viral presence in maternal and placental tissues. Together, these findings advance our understanding of MAYV infection, highlight gestational timing as a factor associated with fetal exposure and suggest the possibility of transmission through routes other than mosquitoes, providing a framework for future studies on this emerging virus.

## Introduction

Mayaro virus (MAYV), an emerging arbovirus of the *Alphavirus* genus and *Togaviridae* family, was first identified in Trinidad and Tobago in 1954 and has since been responsible for sporadic outbreaks across Central and South America. Imported cases have also been reported in Europe and North America, raising concerns at the international public health level as reflected in the epidemiological alert issued by the Pan American Health Organization / World Health Organization in May 2019, which emphasized the need to strengthen epidemiological surveillance and laboratory diagnostic capacity (1). The virus has already been detected in Brazil, Peru, Venezuela, Suriname, Colombia, Ecuador, Bolivia and Panama, with serological evidence also reported in Costa Rica, Guatemala, Mexico and Haiti (2–5). Multiple anthropogenic factors—including climate change, urbanization, human mobility and the expanding distribution of mosquito vectors such as *Aedes aegypti* and *Aedes albopictus*—contribute to the increasing risk of MAYV geographic expansion, even in countries without active transmission. A recent study estimated that 58.9 million people in Latin America reside in ecologically suitable areas for MAYV transmission (6, 7). From a genomic perspective, MAYV possesses a positive-sense single-stranded RNA genome of approximately 11.5 kb, which encodes both non-structural (nsP1–nsP4) and structural proteins (CP, E3, E2, TF, 6k and E1) through two open reading frames (ORFs). The virion is enveloped in a lipid membrane containing E1 and E2 glycoproteins and exhibits high genetic plasticity, which facilitates its adaptation to a broad range of hosts. The viral replication cycle begins with the fusion of the viral envelope to the host cell membrane, followed by translation of viral proteins and the assembly of new virions. MAYV is classified into three genotypes: D (dispersed), L (limited) and N (new), the latter identified in Peru (8–13). In addition to its sylvatic cycle, MAYV is proposed to establish urban transmission through vectors such as *Aedes spp.*, *Culex spp.* and *Sabethes spp.*, with potential vertebrate reservoirs including marsupials, rodents, birds and alligators (14–16).

Clinically, Mayaro disease is generally a self-limiting illness characterized by fever, rash, headache, photophobia, myalgia and persistent polyarthralgia, which may last for months or even years, significantly impacting patients’ quality of life. Hemorrhagic and neurological complications, as well as myocarditis, have also been reported (17). The clinical similarity to other arboviral diseases, such as Dengue and Chikungunya, combined with limited diagnostic capacity in endemic regions, hinders accurate differential diagnosis and contributes to underestimation of the actual disease burden (18). Beyond its typical acute presentation, increasing evidence from other arboviral infections, including Zika and Chikungunya viruses (ZIKV; CHIKV), indicates that clinical outcomes and transmission dynamics may extend beyond the classical vector-borne paradigm, raising questions about the potential involvement of vulnerable populations such as pregnant women and neonates (19–22). Although no specific antiviral treatments or licensed vaccines are currently available, several vaccine candidates are under development, with sustained epidemiological surveillance remaining essential to anticipate and mitigate future outbreaks (3, 8, 23).

Animal models, particularly murine species, have been instrumental in elucidating the pathogenesis of MAYV and in the preclinical evaluation of therapeutic and vaccine candidates. These models faithfully reproduce hallmark clinical manifestations—including ruffled fur, altered gait and ocular irritation—as well as key histopathological features such as arthritis, myositis and tenosynovitis (24, 25). Disease outcomes vary depending on host age and immune status (26–28). While the musculoskeletal system represents the site of the most evident clinical damage, the liver and spleen act as major systemic sites of viral replication, functioning as central hubs for Mayaro virus amplification and dissemination (9, 27). The development of experimental platforms that enable robust and reproducible antiviral testing is essential for advancing MAYV research. In this context, the integration of *in vivo* imaging technologies constitutes a major methodological advance, as it allows real-time, non-invasive monitoring of viral dissemination dynamics using tagged viruses (26, 29).

Here, we establish and refine a murine model as a preclinical platform for the future evaluation of antiviral drugs and therapeutic interventions against MAYV. To validate early systemic dissemination under controlled conditions, we employed a nanoluciferase-expressing MAYV reporter for longitudinal *in vivo* imaging. Complementary experiments using wild-type MAYV were performed to characterize tissue tropism, viral load, hematological and biochemical alterations and placental transcriptional responses following controlled infection. We further assessed gestational stage–dependent placental permissiveness and explored the biological plausibility of vertical and sexual transmission. We hypothesize that gestational timing modulates placental susceptibility to MAYV, defining critical windows of maternal–fetal vulnerability. Together, this integrated framework provides a basis for mechanistic investigation of systemic dissemination, reproductive involvement and maternal–fetal risk during MAYV infection.

## Materials and methods

### Ethics statement

All procedures and experiments were conducted in accordance with the Ethics Committee on Animal Use of the Institut Pasteur de Montevideo (CEUAIP #010-22, in compliance with National Law 18.611 “Procedures for the Use of Animals in Experimentation, Teaching and Scientific Research Activities”) and strictly followed ARRIVE guidelines.

### MAYV infective clones

This study employed wild-type MAYV and nano Luc-expressing MAYV (MAYV-nLuc) infectious clones based on the TRVL 4675 strain, a genotype D virus originally isolated from a human case in Trinidad in the 1950s and kindly provided by Dr. James Weger-Lucarelli (29–31).

### Cell culture

Vero cells were maintained in Dulbecco’s Modified Eagle Medium (DMEM) supplemented with 10% fetal bovine serum (FBS), penicillin (100 U/mL) and streptomycin (100 µg/mL) and maintained at 37 °C, with 5% CO₂.

### Virus generation

Infectious virus was rescued from both WT and MAYV-nLuc infectious clones by transfection of Vero cells using Lipofectamine™ 3000 (Invitrogen), according to the manufacturer’s instructions. Viral supernatants were collected when transfected cells exhibited ≈80% cytopathic effect (CPE) and stored at-80°C as passage P0.

Working viral stocks were generated by sequential infection of Vero cells to produce P1 and P2 passages. The P2 viral stocks were titrated by plaque assay, verified by sequencing and stored at-80°C until further use.

### Mice

BALB/cJ mice (#000651) were bred and maintained at the Laboratory Animal Biotechnology Unit’s animal facility of Institut Pasteur de Montevideo. Mice were housed in IVC Isolator Biocontainment unit (ISOCAGE-N, Tecniplast, Milan, Italy) and acclimatized for one week prior to the experimental procedures. The cages were prepared with wood chip bedding (Toplit 6, SAFE, France), plastic tunnels as enrichment and paper napkins as nesting material. Environmental conditions were maintained at 20 ± 1°C, with 30–70% relative humidity and a 14/10-hour light/dark cycle. The mice were provided with *ad libitum* autoclaved standard diet (D113, SAFE) and filtered autoclaved water. Mice were specific pathogen-free (SPF) certified by IDEXX (USA) quarterly, in accordance with FELASA standards.

### Animal experimental design

Intraperitoneal (i.p.) inoculation was selected to enable controlled and synchronized assessment of early systemic dissemination of Mayaro virus under standardized experimental conditions. Although this route does not reflect natural mosquito-borne transmission, it enables precise dose administration and reproducible systemic exposure, thereby facilitating analysis of early viral dissemination dynamics. A nanoluciferase-expressing MAYV reporter (MAYV-nLuc) was employed for non-invasive *in vivo* imaging. This reporter-based approach was used to define spatial and temporal dissemination patterns. Complementary experiments using wild-type MAYV were subsequently performed to quantify systemic infectious viral loads, characterize hematological and biochemical alterations and assess placental transcriptional responses following controlled infection. Viral doses were optimized according to the specific objectives of reporter-based imaging and pathogenesis studies. Sex-specific analyses were incorporated where biologically relevant, including assessment of reproductive tissue tropism and transmission experiments. This integrated experimental framework enabled systematic evaluation of viral dissemination, reproductive involvement and gestational stage–dependent placental permissiveness (Fig 1).

**Figure 1.**
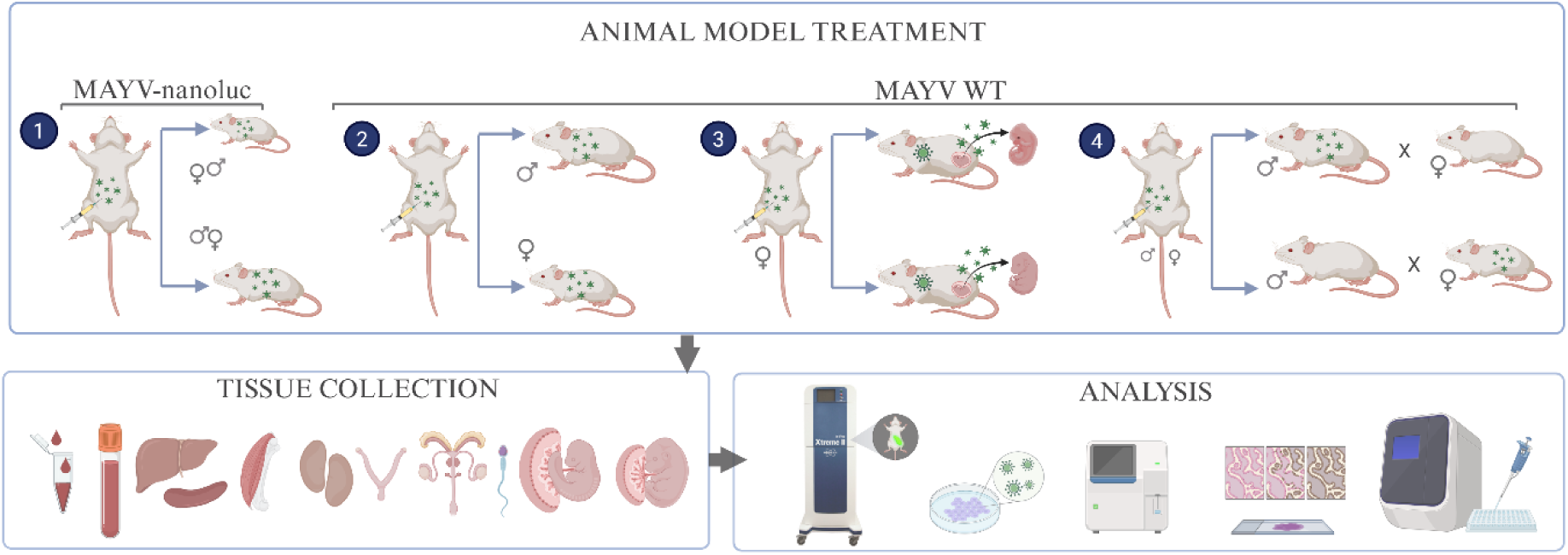
Experimental design for longitudinal assessment of early systemic dissemination of Mayaro virus in mice. Mice were i.p. inoculated with a bioluminescent Mayaro virus to enable longitudinal, non-invasive monitoring of early viral dissemination by *in vivo* imaging. Parallel groups were infected with wild-type Mayaro virus to evaluate tissue tropism and transmission-related outcomes. Blood and lymphoid, visceral, musculoskeletal and reproductive tissues were collected for downstream analysis.

### *In vivo* imaging evaluation with MAYV-nLuc infection

*In vivo* bioluminescence imaging frequency was performed over a 10-day observation window to capture the peak of viral replication and the onset of immune responses associated with recovery from acute infection, based on previous reports (24, 25, 32). Age groups were selected according to their susceptibility to MAYV infection, with younger animals exhibiting higher susceptibility (33). For this study, four-and eight-week-old BALB/cJ female mice were i.p. infected with 1 x 10^4^ PFU of MAYV-nLuc resuspended in 100 μL PBS (n=4 per age group). Mock-infected animals received PBS only (n = 2; one animal per age group). The viral dose was selected based on pilot optimization experiments to ensure robust reporter detection while maintaining animal welfare. To perform *in vivo* luminescence imaging, the animals were i.p. injected with 20 µg of Furimazine (TargetMol®, MA, USA) in 100 µL of PBS, anesthetized with isoflurane (3–4% for induction and 1-2 % for maintenance) and imaged at defined time points using an In-Vivo Xtreme II™ system (Bruker Daltonik GmbH, Billerica, MA, USA). Image acquisition and analyses were performed using MolecularImaging software V.7.5.3.22464 (Bruker Daltonik Gmbh, Billerica, MA, USA). Thresholds and exposure times were kept constant across time points for longitudinal comparability. Bioluminescent signal reflects reporter expression and may not directly correlate with infectious viral titers. Imaging was performed at 1-, 2-, 6-and 10-day post infection (dpi). Animals were necropsied at 6 dpi (n=2 per age group) and at 10 dpi (remaining animals). Organs collected for confirmation of infectious viral particles included the liver, kidneys, spleen and *quadriceps femoris* muscle. Peripheral tissues exhibiting luminescent signal—tail, ear, forelimb or hindlimb—were also sampled.

### Viral tropism of Mayaro WT and health status

Twelve BALB/cJ female and twelve BALB/cJ male mice (4 weeks old) were i.p. infected with 1 x 10⁵ PFU of MAYV WT resuspended in 100 μL of PBS (n=24). Mock-infected mice received the PBS diluent alone (n=4). The viral dose was selected based on prior pilot experiments as the lowest dose that consistently allowed detection of infectious viral particles across tissues during acute infection. Blood samples were collected pre and post-infection for hematological and biochemical parameter analyses at 0-, 24-and 48-hour post infection (hpi). At 24 hpi, six animals of each sex were euthanized under deep anesthesia induced by ketamine-xylazine mixture administered via the i.p. route (ketamine 110 mg/kg; Pharmaservice, Ripoll Vet, Montevideo, Uruguay; xylazine 13 mg/kg; Seton 2%; Calier, Montevideo, Uruguay), followed by cervical dislocation. Organs collected for quantification of infectious viral particles included the liver, kidneys, spleen, *quadriceps femoris* muscle, ovaries, uterus, seminal vesicles and testes. The same procedures were performed on the remaining animals at 48 hpi.

### MAYV WT vertical transmission

Seventeen BALB/cJ female mice (8-weeks old) were synchronized in their estrous cycle by exposing them to male dirty bedding for 72 h prior to mating (34). Mating was then performed by placing one proven male in a cage with two females for 48 h and the presence of a vaginal plug was checked daily from the following morning as a positive sign of mating to be considered day 0.5 of pregnancy. At 7.5-8.5-day post-coitum (dpc), six females were i.p. infected with 1 x 10⁵ PFU of MAYV WT resuspended in 100 μL of PBS, while two mock-infected received only PBS diluent (n = 6 infected; n = 2 mock). The females were euthanized and necropsied at 48 hpi (9.5-10.5 dpc). Blood samples were collected prior to euthanasia for post-infection hematological and biochemical analyses. Organs collected for quantification of infectious viral particles and histological analysis included the liver, kidneys, *quadriceps femoris* muscle, ovaries, uterus, placenta and fetus. The same procedures were performed on the remaining pregnant females infected at 13.5–14.5 dpc (n = 7 infected; n = 2 mock), which were necropsied at 48 hpi (15.5-16.5 dpc).

### MAYV WT sexual transmission

To evaluate sexual transmission in both directions, BALB/cJ mice were mated in a 1:1 ratio under controlled conditions. For female-to-male transmission, seven 8-week-old females were synchronized in their estrous cycles by exposure to male dirty bedding for 72 h prior to mating, as described above and were infected i.p. with 1 × 10⁵ PFU of MAYV in 100 μL of PBS 12 h before pairing with uninfected males. For male-to-female transmission, six synchronized 8-week-old females were paired with males infected i.p. with the same viral dose and schedule. Infected animals were necropsied at 48 hpi, whereas their uninfected partners at 48 h after mating. The liver, spleen, *quadriceps femoris* muscle, ovaries, uterus, seminal vesicles and testes were collected for infectious viral particle detection by plaque assay and for RT-qPCR analysis in organs from exposed animals. Sperm from the bilateral cauda epididymides was collected for plaque assay and RT-qPCR analysis. For analytical purposes, an exposed animal was defined as an uninfected individual mated with an infected partner.

### Hematology and biochemistry profile

For hematological analysis, 20 µL of blood per animal were collected and stored in 0.5 mL microtubes containing potassium EDTA salts (W anticoagulant, Wiener Lab, Rosario, Argentina) at a 1:10 ratio (EDTA: blood). Samples were analyzed using an Auto Hematology Analyzer BC-5000 Vet (Mindray, Shenzhen, China). The parameters evaluated included white blood cells (WBC), neutrophils (Neu), lymphocytes (Lym), monocytes (Mon), eosinophils (Eos), basophils (Bas), red blood cells (RBC), hemoglobin (HGB), hematocrit (HCT), mean corpuscular volume (MCV), mean corpuscular hemoglobin (MCH), mean corpuscular hemoglobin concentration (MCHC) and platelets (PLT).

For biochemical blood profile analysis, 100 μL of whole blood per animal was collected and mixed with 0.5 μL of sodium heparin (Heparin Na, 20 U/mL). Using the Chemistry Analyzer Pointcare® V3 (Tianjin MNCHIP Technologies Co. Ltd, Tianjin, China), the following parameters were evaluated: total protein (TP), albumin (ALB), globulin (GLO), total bilirubin (Tbil), alanine aminotransferase (ALT), aspartate aminotransferase (AST), gamma-glutamyl transferase (GGT), blood urea nitrogen (BUN) and creatinine (CRE).

### Tissue collection and processing

Samples collected from each organ of interest were weighed, rapidly frozen in liquid nitrogen under a biosafety cabinet and stored at-80°C until processing. Samples were homogenized using zirconium beads (0.5 mm/tube) and 800 µL of sterile 1× DPBS in a Precellys® 24 homogenizer (Bertin Technologies, France), (2 min, power 10). Homogenates were clarified by centrifugation (6,000 × g, 5 min) and supernatants were stored at −80 °C for viral titration or RT-qPCR analysis.

### Viral titration

For viral titration, confluent Vero cell monolayers in 12-well plates (2.6 × 10⁶ cells per plate) were inoculated with tissue homogenate–derived supernatants and their 10-fold serial dilutions (200 µL per well) and incubated at 37 °C with 5% CO₂ for 1 h, with gentle rocking every 15 min to ensure uniform inoculum distribution. Mock-infected wells served as negative controls. After adsorption, cells were overlaid with DMEM containing 0.8% agarose, 1.6% FBS, penicillin and streptomycin and incubated for 48 h. Monolayers were then fixed with 4% formaldehyde, stained with crystal violet and plaques were counted to determine viral titers, expressed as plaque-forming units per gram of tissue (PFU/g).

### Mouse sperm collection

The bilateral cauda epididymides were collected in a 60 mm plastic Petri dish and incised with several cuts using fine scissors, releasing the sperm into a 50 µL drop of Dulbecco’s Phosphate Buffered Saline 1x (DPBS, Sigma-Aldrich) and stored at −80°C until further processing. Later, sperm samples were thawed and resuspended in 150µL of PBS. Zirconium beads were added to each sample and disruption was performed using a Bullet Blender® STORM 24 (Next Advance Inc., New York, USA) for one minute at speed 8. After centrifugation, 100 μL of supernatant was divided into two aliquots. To assess the presence of MAYV, one aliquot was subjected to cell culture amplification followed by plaque assay, whereas the other was analyzed directly for viral RNA by RT-qPCR.

### Viral RNA extraction and detection

Viral RNA from tissues was extracted from clarified tissue homogenate supernatants, as described above, using the QIAamp® Viral RNA Mini Kit (QIAGEN), according to the manufacturer’s instructions. RNA from sperm samples was extracted from TRIzol-treated homogenates using the Direct-zol RNA Miniprep Kit (Zymo Research), following the manufacturer’s guidelines. RNA samples were eluted in 25µL of nuclease-free water and stored at-80°C until further use (35). Detection was performed using a one-step real-time reverse transcription PCR (RT-qPCR) assay, employing the primers and probe listed in Table 1, which target an 86-bp region of the MAYV nsP1 gene. Reactions were carried out using TaqMan™ Fast Virus 1-Step Master Mix (Thermo Fisher Scientific) on a QuantStudio™ 7 Pro Real-Time PCR System (Applied Biosystems). Due to limited sample availability, sperm-derived RNA samples were analyzed in technical duplicates. The assay was used for qualitative detection (presence/absence) of viral RNA.

**Table 1:**
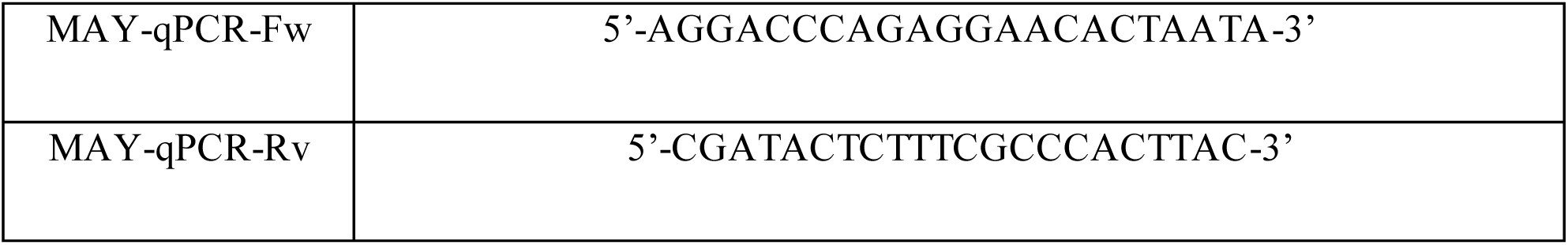
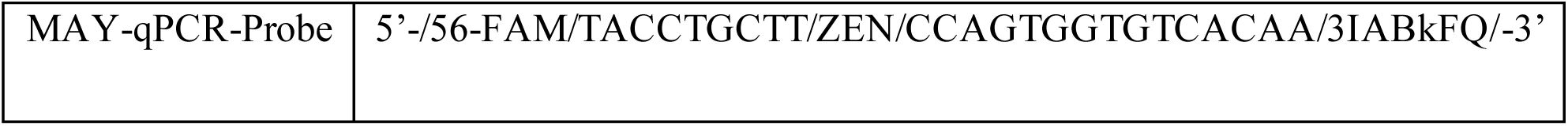
primers and probe used for MAYV detection by qPCR.

### Cell culture–based isolation of virus from sperm samples

Due to the limited volume of sperm samples, processed samples were subjected to passage in Vero cells to allow potential biological amplification prior to titration. Vero cells were seeded in 96-well plates at 5,000 cells per well in 100 µL of DMEM supplemented with 5% FBS, penicillin (100 U/mL) and streptomycin (100 µg/mL). The following day, cells were inoculated with 100 µL of processed sperm samples. Mock-infected wells receiving culture DMEM alone served as negative controls. After 72 h of incubation at 37 °C, supernatants were harvested and the presence of infectious viral particles was assessed by plaque assay. Cytopathic effects were monitored using inverted optic microscope Zeiss Primovert, Carl Zeiss Suzhou Co., LTD, China.

### Transcriptomics in placenta

For the RNA-seq experiment, three placentas per condition (control uninfected; infected at 7.5-8.5 dpc and infected at 13.5-14.5 dpc) were selected and processed for total RNA extraction. Placentas stored at −80 °C were disrupted in TRI Reagent using a Bullet Blender® STORM 24 (Next Advance Inc., New York, USA) with 0.5 mm zirconium oxide beads (ZROB05, Next Advance Inc.) for 10 min at maximum speed. Following homogenization, the samples were centrifuged to remove beads and debris. The upper aqueous phase was collected and purified using the Direct-zol RNA kit (Zymo Research, USA) according to the manufacturer’s instructions. RNA was quantified with RNA HS Qubit (Invitrogen, USA). RNA quality assessment, library construction and sequencing were performed by Macrogen (Seoul, South Korea). Libraries were prepared using TruSeq stranded total RNA with Ribo-Zero Gold kit (Illumina, USA). Sequencing was performed on an Illumina NovaSeq X platform. Paired-end reads of 100 base pairs were obtained.

Raw paired-end RNA-seq reads were aligned to the *Mus musculus* reference genome (GRCm39, Ensembl release 114) using STAR v2.7.8a. Aligned reads were quantified at the gene level using featureCounts v2.0.2 (Subread package). The Ensembl GTF annotation file (Mus_musculus.GRCm39.114.gtf) was used.

Downstream analysis was conducted in R v4.3.0 using the DESeq2 package. The raw count matrix was filtered to retain genes with a minimum of 10 total reads across all samples. A DESeq2 dataset was constructed with experimental conditions as the main design factor. Variance stabilizing transformation (VST) was applied to normalize data prior to visualization and clustering. Differential expression analyses were performed for multiple contrasts of interest and log2 fold changes were shrinkage-corrected using the “normal” method of lfcShrink. Genes with an adjusted p-value <0.05 were considered significantly differentially expressed.

Volcano plots were generated with ggplot2, where significant genes were highlighted (adjusted p <0.05). Heatmaps of differentially expressed genes were created with pheatmap R package, using variance-stabilized expression values scaled by gene (row-wise z-scores). Color schemes were defined with the RColorBrewer package. Additional hierarchical clustering of differentially expressed genes was performed to assess global expression patterns across experimental groups. Gene ontology analysis was performed using Metascape (36).

### Immunohistochemistry

Immediately after the necropsy, placenta and fetus from mid-stage of pregnancy were fixed in 10% neutral buffered formalin (pH 7.4) for further processing. The samples were then decalcified by immersion and agitation in a 3% aqueous nitric acid solution for 24 h. Once decalcification was complete, histological processing for paraffin embedding began. Once the paraffin blocks were obtained from each sample, successive 4µm-thick sections were made with a manual rotary microtome (Slee, Germany) and the successive sections were mounted on slides and prepared for immunohistochemistry against MAYV immunoreaction, using a rabbit polyclonal primary antibody (dilution 1:100) according to a previous immunofluorescence protocol developed for immunocytochemistry in cell cultures (37, 38). Histological and immunohistochemical blinded examinations were performed using transmitted light microscope (Nikon Eclipse E200^®^, Nikon Corporation, China) by two different veterinary pathologists.

## Data Analysis

Statistical analyses were performed using GraphPad Prism version 10 (GraphPad Software, San Diego, CA, USA). Viral load data (PFU/g) were log₁₀-transformed prior to analysis. Values below the limit of detection (LOD) were imputed as LOD/2 (630 PFU/g) before transformation. Viral load data were analyzed using mixed-effects models (REML), including time point, tissue, sex and their interactions as fixed effects and individual animal as a random effect to account for within-subject variability. Post hoc comparisons were performed using Tukey’s multiple comparisons test. Hematological and biochemical parameters with repeated measurements were analyzed using mixed - effects models (REML), including time, infection status and their interaction as fixed effects and individual animal as a random effect to account for within-subject correlation. For cross-sectional analyses without repeated measures involving two factors (infection status and gestational stage at 48 hpi), two-way ANOVA followed by Sidak’s multiple comparisons test was used. Model assumptions were evaluated by assessing normality of residuals using the Shapiro–Wilk test and by visual inspection of residual plots. RNA-seq differential expression analysis was performed using DESeq2. Genes with fewer than 10 total counts across all samples were filtered out. Variance stabilizing transformation (VST) was applied prior to visualization and clustering. Differentially expressed genes were identified using adjusted p-values (Benjamini–Hochberg correction), with a significance threshold of adjusted p < 0.05. Data are presented as individual values with median lines for visualization, whereas statistical inference is based on model-derived estimates. All statistical tests were two-tailed and p < 0.05 was considered statistically significant.

## Results

### Bioluminescence reveals rapid dissemination and longitudinal persistence of MAYV-nLuc signal

Luminescence was first detected at 24 hpi (1 dpi) and remained detectable through the predetermined endpoint at 10 dpi. Signals were identified in the forelimbs, hindlimbs, tail, ears and abdominal region. The abdominal signal decreased after 6 dpi, whereas residual luminescence persisted in peripheral sites, remaining detectable mainly in the limbs at 10 dpi (Fig 2).

**Figure 2.**
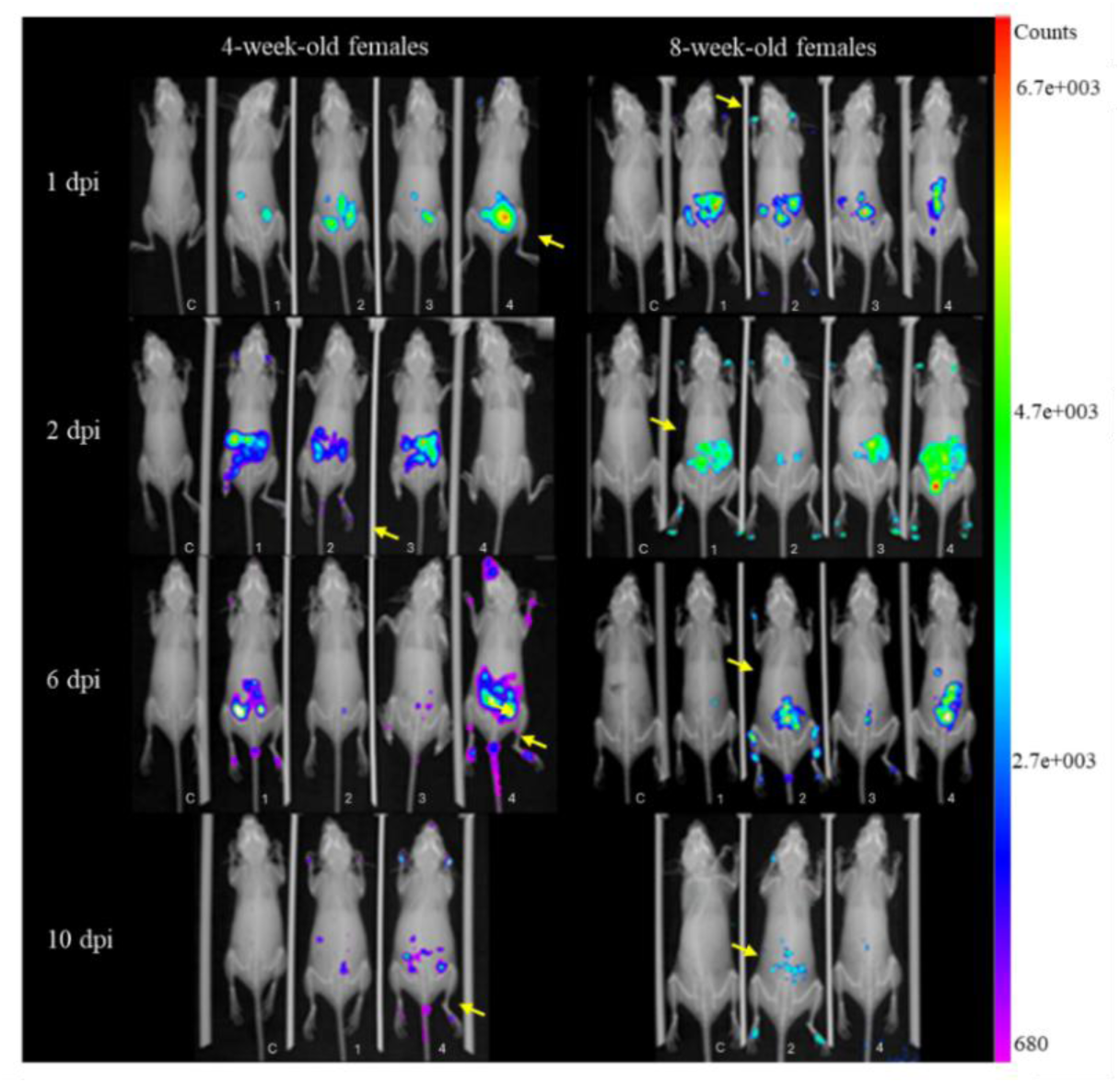
*In vivo* bioluminescence imaging of MAYV-nLuc infection in females at 4 and 8 weeks-old following i.p. inoculation. (representative image). Luminescent signals (yellow arrows) were detected in the limbs, tail, ears and abdominal region at 1 (24 hpi), 2 (48 hpi), 6 and 10 dpi. Abdominal signals declined after 6 dpi, whereas residual luminescence persisted mainly in the limbs at later time points. Images from all animals in each age group are shown. All images were acquired and displayed using identical exposure settings and a fixed color scale to enable direct comparisons over time. C, mock-infected control.

To assess the presence of infectious virus, plaque assays were performed on tissues collected at 6 and 10 dpi. Visceral organs (liver, spleen, muscle and kidneys) from both 4-and 8-week-old mice were analyzed, along with peripheral tissues (limbs, tail and ear) displaying detectable bioluminescent signal. In 4-week-old animals, infectious viral particles were recovered exclusively from peripheral tissues exhibiting luminescence including limb, tail and ear samples, whereas all visceral organs remained negative. Viral loads in these peripheral compartments ranged from 1.49×10³ to 2.98×10⁵ PFU/g, with the highest titers detected in tail and limb tissues. In contrast, no infectious viral particles were detected in any tissues from 8-week-old animals, including peripheral tissues exhibiting bioluminescent signal.

### Temporal dynamics and tissue tropism of MAYV infection in 4-week-old mice

Viral titers exhibited time-and tissue-dependent differences during acute infection (Fig 3A). Mixed-effects analysis with Tukey’s multiple comparisons identified a significant increase in seminal vesicles at 48 hpi compared with 24 hpi (Mean difference = 0.9452, SE of difference = 0.3397, *p* = 0.0284). Model-based estimates are reported as mean differences (± SE), while graphical representations show individual values with median lines due to non-normal data distribution. At 24 hpi, no significant inter-tissue differences were detected, consistent with a relatively homogeneous early dissemination pattern. By 48 hpi, tissue-specific differences emerged, with spleen titers exceeding those in liver (Mean difference = 0.8391, SE of difference = 0.2150, *p* = 0.0352) and muscle (Mean difference = 1.257, SE of difference = 0.2840, *p* = 0.0156). This substantial difference suggests the spleen as a primary site of viral accumulation compared to muscular tissue.

**Figure 3.**
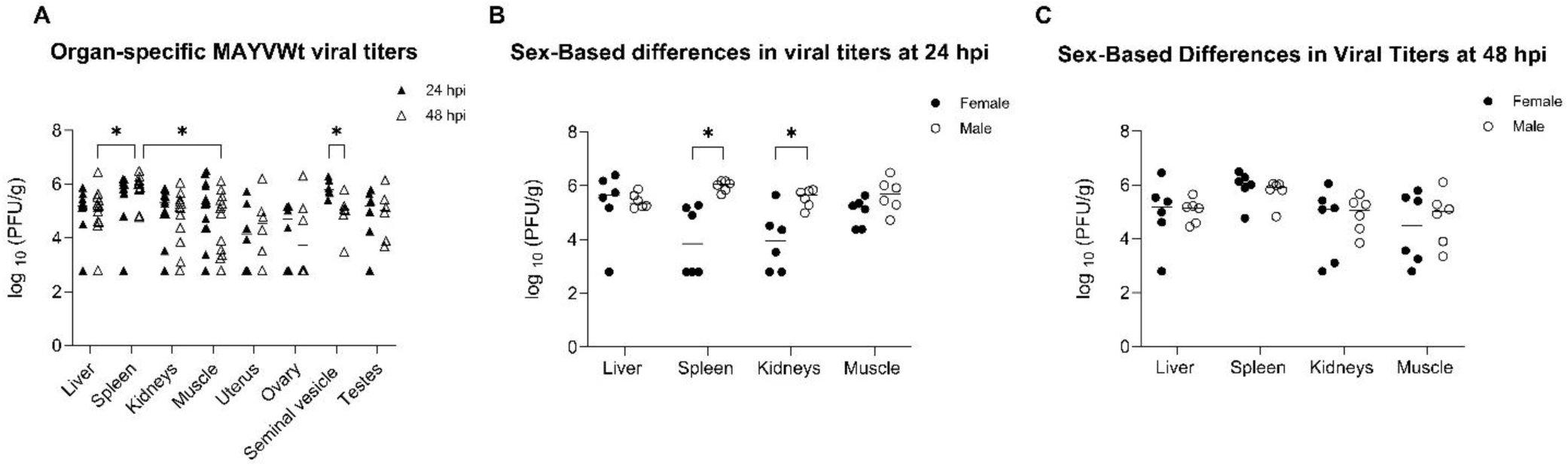
Early systemic dissemination with time-dependent tissue tropism and transient sex effects. Viral loads (log₁₀ PFU/g) across tissues at 24 and 48 hours post-infection (hpi) and sex-based comparisons at each time point. (A) Viral loads across tissues at 24 and 48 hpi. (B) Sex-based comparison of viral loads across tissues at 24 hpi. (C) Sex-based comparison at 48 hpi. Viral loads were analyzed using mixed-effects models (REML) with time point and tissue as fixed effects and individual animal as a random effect, followed by Tukey’s multiple comparisons test. Values below the limit of detection were assigned a value of LOD/2 (630 PFU/g) prior to transformation. Each dot represents an individual animal; horizontal lines indicate median values. Only statistically significant comparisons are indicated. Statistical significance was defined as p < 0.05 (*), p < 0.01 (**) and p < 0.001 (***).

Sex-based analyses revealed transient differences at 24 hpi (Fig 3B). Specifically, significant female–male differences were observed in the spleen (mean difference = 1.593, SE of difference = 0.5278, *p* = 0.0109) and kidneys (mean difference = 2.038, SE of difference = 0.4765, *p* = 0.0159). In both tissues, males exhibited substantially higher viral titers than females at this time point. Organ-specific viral distribution also varied within each sex at 24 hpi. Among females, viral loads in the kidneys were significantly higher than in the muscle (mean difference = 1.078, SE of difference = 0.2911, *p* = 0.0493). Conversely, in males, the spleen showed a significantly greater viral burden compared to the liver (mean difference = 0.5601, SE of difference = 0.1205, *p* = 0.0205). By 48 hpi, no significant differences in viral loads were observed either between sexes or among tissues within each sex (Fig 3C). This lack of statistical significance (*p* > 0.05 for all comparisons) indicates that the early sex-dependent effects and tissue-specific disparities observed at 24 hpi are not sustained as the infection progresses toward later stages.

### Early hematological alterations during acute MAYV infection

Hematological analysis revealed alterations in leukocyte profiles during acute infection. Total white blood cell counts (WBC) decreased trend at 24 and 48 hpi, accompanied by increased neutrophil (NEU) and decreased lymphocyte (LYM) percentages (Fig 4A –C). Platelet counts (PLT) showed a decreased at both time points although variability across animals was observed (Fig 4D). Eosinophils (EOS) decreased, basophils (BAS) increased at 48 hpi and monocytes (MON) remained stable in infected animals but decreased in mock controls (Fig S1A–C). Red blood cell counts (RBC) increased at 48 hpi, while hemoglobin (HGB) and hematocrit (HCT) increased at both time points (Fig S1D –F). Mean corpuscular volume (MCV) increased at 24 and 48 hpi, mean corpuscular hemoglobin (MCH) increased at 24 hpi and mean corpuscular hemoglobin concentration (MCHC) remained unchanged (Fig S3A–C). Red cell distribution width (RDW-CV and RDW-SD) decreased at both time points. Mean platelet volume (MPV) increased, while platelet distribution width (PDW) decreased at 48 hpi. Procalcitonin (PCT) decreased in infected animals without changes in mock controls (Fig S3D–H). Statistically significant differences are indicated in the corresponding figures.

**Figure 4.**
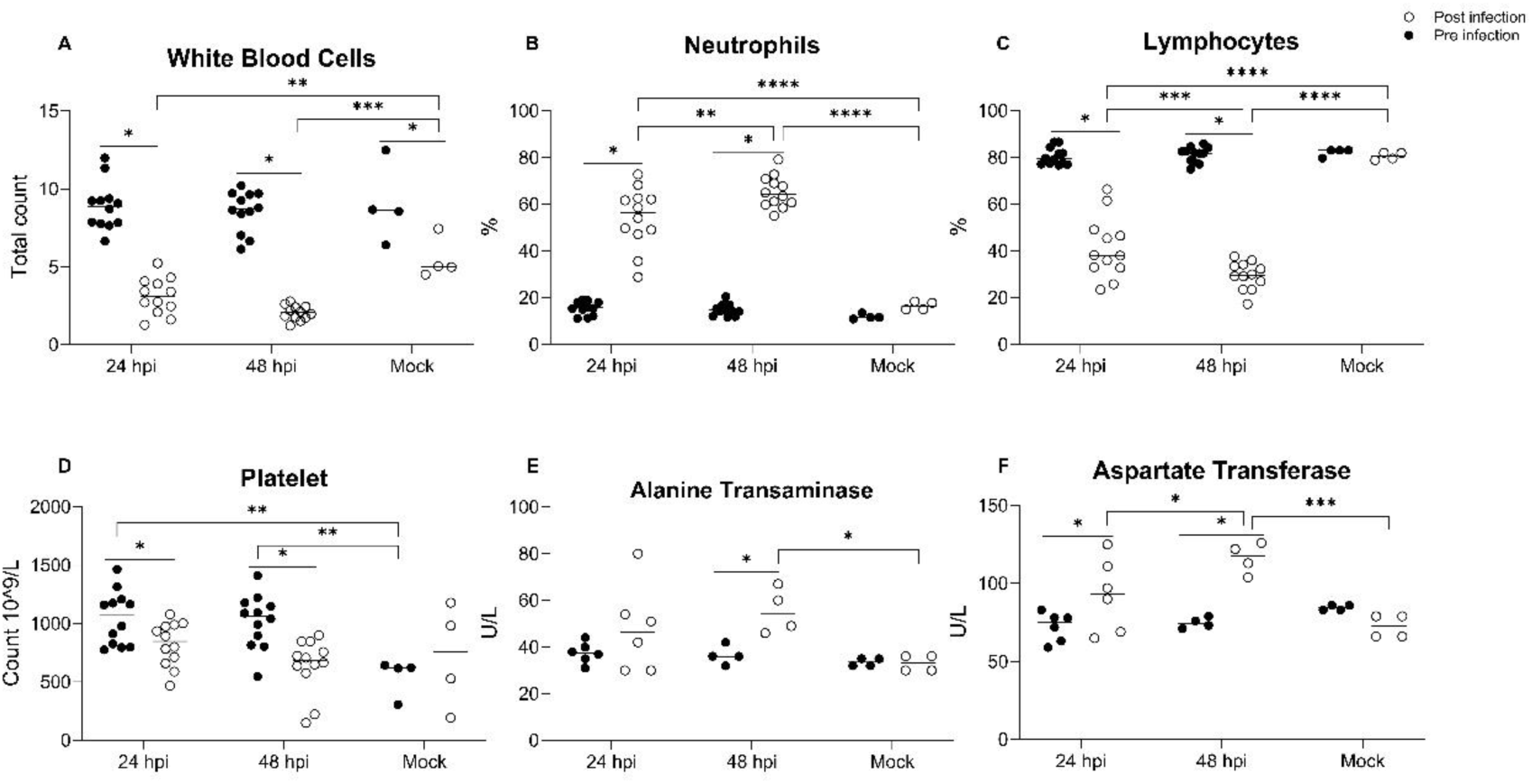
Early hematological and hepatic changes following acute MAYV infection. Hematological and hepatic parameters during acute MAYV infection at 24 and 48 hours post-infection (hpi) compared with mock controls. (A) Total white blood cell (WBC) count, (B) neutrophil percentage (NEU), (C) lymphocyte percentage (LYM), (D) platelet count (PLT), (E) alanine transaminase (ALT) and (F) aspartate transaminase (AST). Each dot represents an individual animal; horizontal lines indicate median values. Statistical analyses were performed using mixed-effects models (REML) for repeated measures with time and condition as fixed effects and individual animal as a random effect. Differences were considered statistically significant at p < 0.05 (*), p < 0.01 (**) and p < 0.001 (***).

In biochemical and metabolic parameters, we observed that: Aspartate transaminase (AST) at both 24 and 48 hpi and increased alanine transaminase (ALT) at 48 hpi (Fig 4E–F). Creatinine (CRE) remained stable, whereas blood urea nitrogen (BUN) decreased at 24 hpi and showed a decreasing trend at 48 hpi, including in mock animals (Fig S1G–H). Total protein (TP) decreased at 48 hpi, while albumin (ALB), globulin (GLO) and gamma-glutamyl transferase (GGT) remained unchanged (Fig S2A–C, E). Total bilirubin (TBIL) decreased at 24 hpi (Fig S2D). Glucose (GLU) increased at both time points and body weight (BW) decreased at 24 and 48 hpi (Fig S2G–H). Statistical significance is indicated in the corresponding figures.

### Gestational stage–dependent differences in MAYV distribution during pregnancy

At 48 hpi, viral titers exhibited significant variations based on gestational stage across multiple tissues (Fig 5). Mixed-effects modeling followed by Tukey’s multiple comparison test revealed significantly higher infectious viral loads at 48 hpi in tissues analyzed at 15.5–16.5 dpc (pregnant mice infected at 13.5-14.5 dpc) compared with those analyzed at 9.5–10.5 dpc (pregnant mice infected at 7.5-8.5 dpc). This stage-dependent increase was observed in the spleen (mean difference = 0.5751, SE of difference = 0.1624, *p* = 0.0213), liver (mean difference = 1.924, SE of difference = 0.1552, *p* = 0.0011), kidneys (mean difference = 1.691, SE of difference = 0.3097, *p* = 0.0055), uterus (mean difference = 1.660, SE of difference = 0.6606, *p* = 0.0403) and placenta (mean difference = 0.7328, SE of difference = 0.2277, *p* = 0.0061). However, infectious virus remained undetectable in fetal tissues at this same stage (15.5–16.5 dpc), with titers falling below the limit of detection. This suggests that even when the placenta is significantly infected, the virus remained undetectable at the fetal compartment in an infectious form. Within early gestation (9.5–10.5 dpc), tissue-specific differences were observed, with the spleen showing higher titers than the liver and kidneys. At mid - gestation, infectious viral particles remained detectable in maternal and placental compartments, supporting ongoing systemic dissemination despite restricted recovery of infectious virus from fetuses.

**Figure 5.**
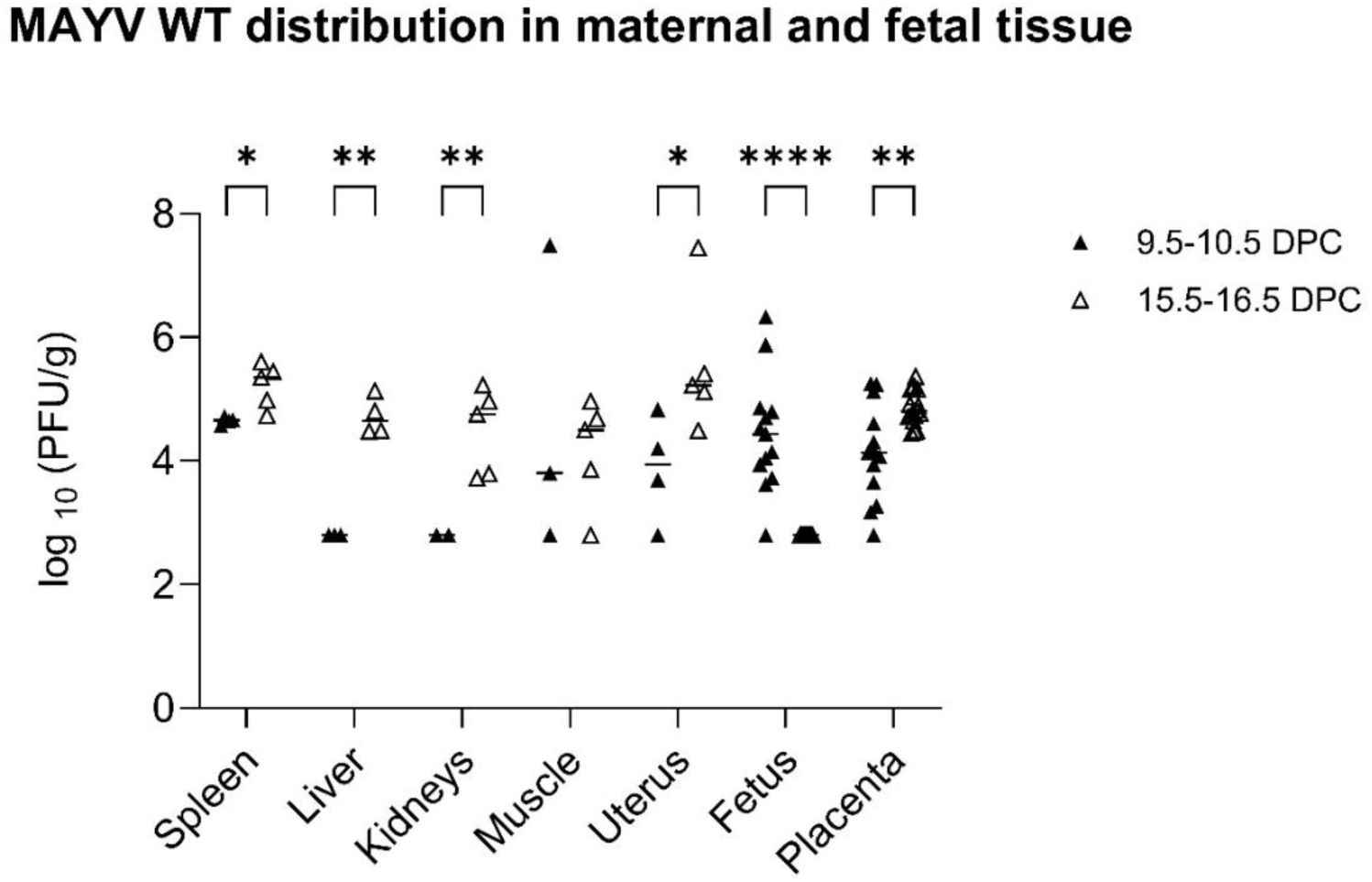
Gestational stage shapes maternal–fetal distribution and restricts recovery of infectious virus from mid-gestation fetuses. Infectious viral loads (log₁₀ PFU/g) in maternal and fetal tissues at 48 hours post-infection (hpi) according to gestational stage. Females were infected at early (7.5 –8.5 dpc; analyzed at 9.5–10.5 dpc) or mid-gestation (13.5–14.5 dpc; analyzed at 15.5–16.5 dpc). Each dot represents an individual animal; horizontal lines indicate median values. Viral loads were analyzed using mixed - effects models (REML) with gestational stage and tissue as fixed effects and individual animal as a random effect, followed by Tukey’s multiple comparisons test. Values below the limit of detection were assigned a value of LOD/2 (630 PFU/g) prior to transformation. Only statistically significant comparisons are indicated. Statistical significance was defined as p < 0.05 (*), p < 0.01 (**) and p < 0.001 (***).

### Exploratory analysis of hematological parameters at 48 hpi revealed alterations in leukocyte profiles across gestational stages

Infected females at both gestational stages exhibited higher neutrophil and lower lymphocyte percentages compared with mock controls (Fig S4A–B). Total white blood cell (WBC) counts were reduced in infected animals (Fig S4C). Platelet (PLT) counts showed variability between groups without a consistent pattern across gestational stages, while red blood cell (RBC) counts and procalcitonin (PCT) levels remained comparable (Fig S4D–F). Given the limited number of mock controls, comparisons involving these groups should be interpreted with caution.

### Immunohistochemistry confirms placental and fetal localization of MAYV at mid stage

Immunohistochemical analysis revealed clear differences in MAYV antigen distribution between infected and non-infected placental and fetal tissues. In placentas from infected females, moderate granular cytoplasmic staining was observed in a substantial proportion of giant trophoblast cells within the maternal–fetal junction zone (basal plate), indicating focal placental infection. In contrast, placentas from non-infected pregnant females showed only nonspecific erythrocyte staining and otherwise negative to mild background labeling. In fetuses from infected females, intense immunostaining was detected across multiple organs, particularly in the intestinal visceral epithelium, developing bone (growth cartilage) and other collagen-rich tissues, including cartilage and dermis. Calcified bone exhibited no staining or only minimal labeling in both intensity and extent. Fetuses from non-infected females were largely negative, showing only nonspecific erythrocyte staining and minimal background labeling in collagen-rich tissues (Fig 6).

**Figure 6.**
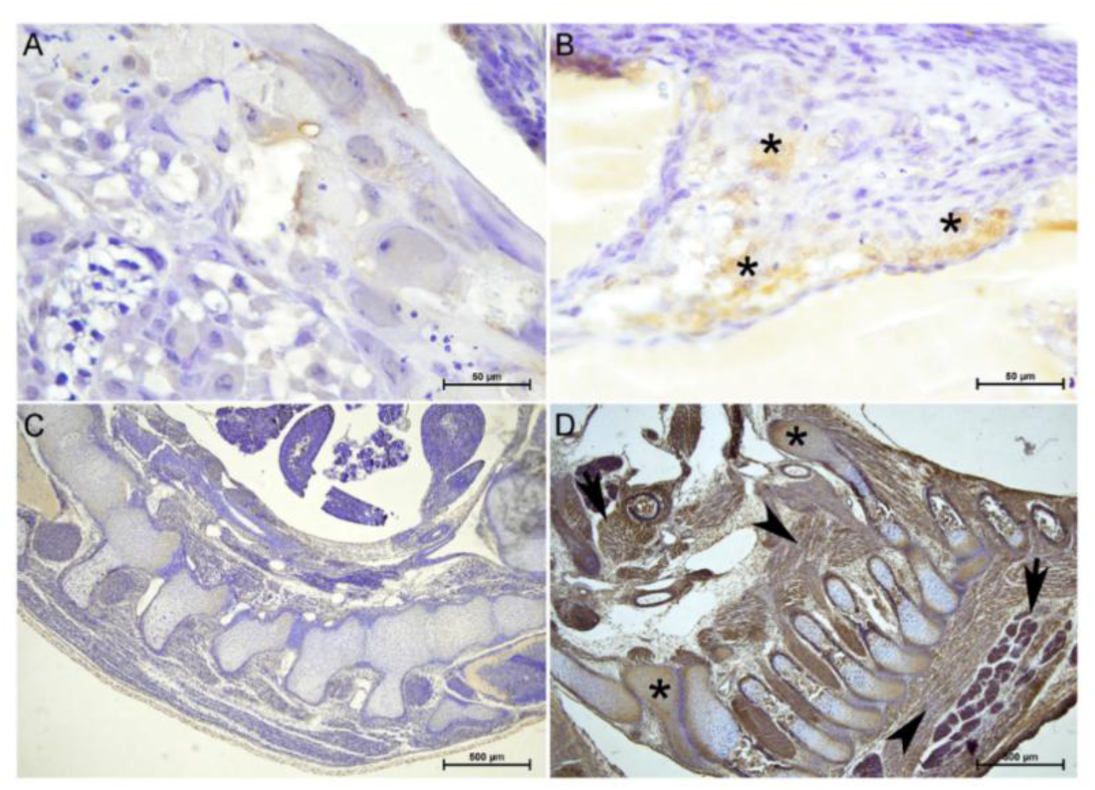
IHC against MAYV. A) Placenta of an uninfected mouse (control): Negative to mild (-/+) immunostaining of isolated trophoblasts (foam giant cells) from the basal plate of the maternal-fetal junction is observed. Scale bar = 50µm, 40×. B) Placenta of an experimentally infected mouse: Mild to moderate, granular intracytoplasmic staining of foci of trophoblasts (foam giant cells) from the basal plate of the maternal-fetal junction is observed (asterisks). Scale bar = 50µm, 40×. C) Fetus of the same uninfected mouse of Figure A: No immunostaining against MAYV is observed (-). Scale bar = 500µm, 4×. D) Fetus of the same experimentally infected mouse of Figure B: There is intense immunostaining in several organs, particularly in uncalcified developing bone and growth cartilage (asterisks), as well as other collagen-rich tissues (cartilage and dermis) (arrowheads) and the perimysium of skeletal muscles (arrows). Already calcified bone shows no staining or minimal staining in intensity and extent. Scale bar = 500µm, 4×.

### Placental transcriptional responses to MAYV are gestational stage-dependent

To investigate placental transcriptional responses to MAYV infection, differential gene expression was performed. The profiles revealed gene expression patterns influenced by viral infection at early and mid-stages: 208 genes were upregulated and 60 downregulated at the early stage and 139 genes were upregulated and 1 downregulated at the mid stage (Fig 7A and Table S1). Among the upregulated genes, 50 were shared between both stages, leaving 158 stage-specific upregulated genes at the early stage and 89 at the mid stage. These data suggest a robust transcriptional response to infection during the early stage of pregnancy, while the response in mid pregnancy, although sharing common pathways, is comparatively attenuated.

**Figure 7.**
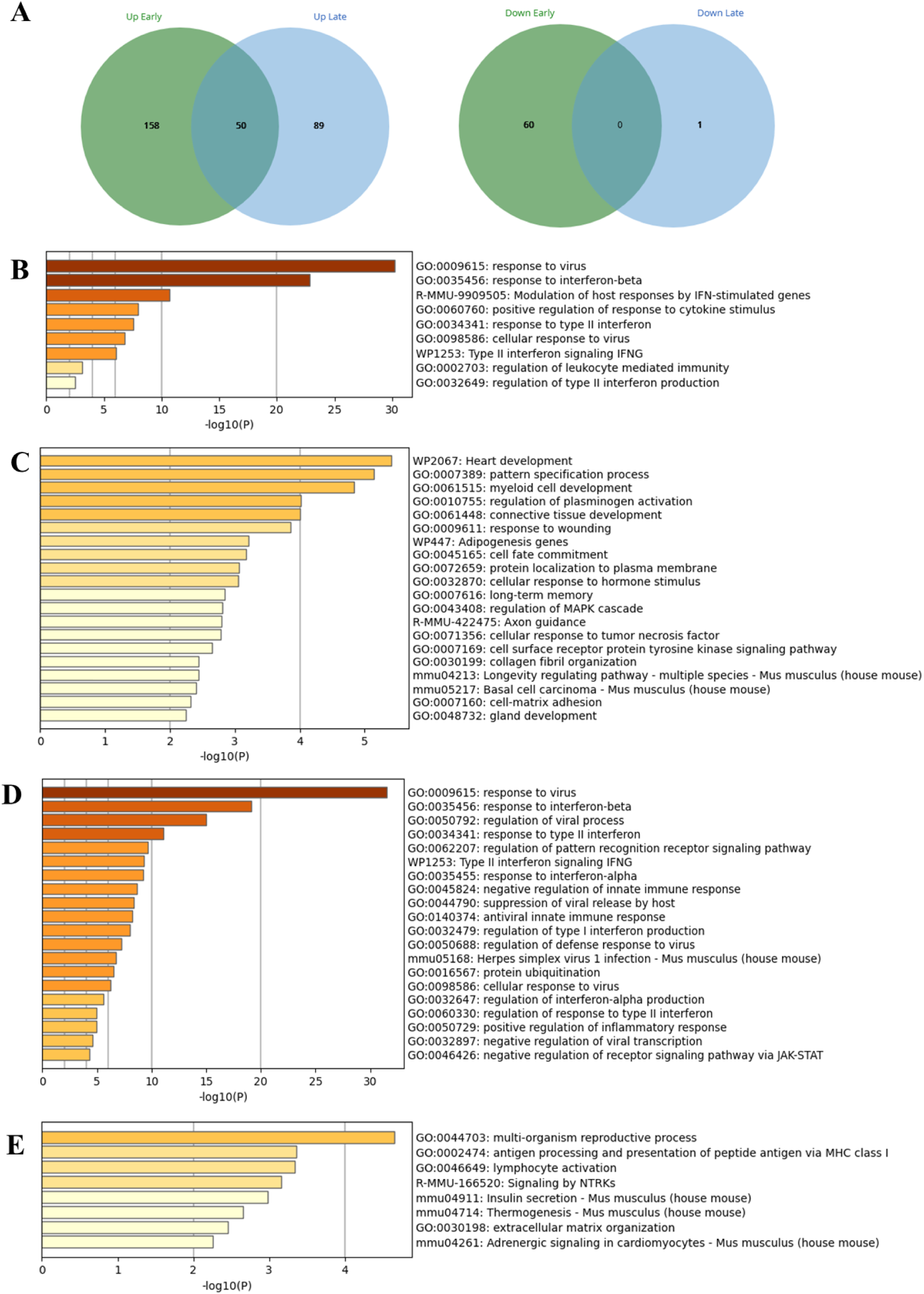
Transcriptomic profiles and Gene Ontology enrichment of placental response to MAYV infection at early and mid-gestational stages. A) Comparison of placentas infected at 7.5–8.5 dpc (early) and 13.5–14.5 dpc (mid) versus controls revealed 208 up-/60 down-regulated genes at early stage and 139 up-/1 down-regulated at mid stage, with 50 genes shared between stages. B) Shared genes were enriched for antiviral and interferon-mediated responses. C) Early-stage-specific genes were associated with immune, developmental and metabolic pathways. D) Mid-stage-specific genes were mainly related to type II interferon and innate immune modulation. E) Early-stage down-regulated genes were enriched for reproductive, metabolic and signaling processes.

Gene Ontology (GO) enrichment analysis of the 50 shared upregulated genes revealed significant association with antiviral defense pathways, including responses to viral infection, interferon-stimulated genes (ISGs) activity and cytokine-mediated signaling (Fig 7B).

The 158 genes uniquely upregulated in the early stage were enriched not only for the immune pathways but also for distinct biological processes, including heart development, myeloid cell development, regulation of plasminogen activation, connective tissue development and adipogenesis (Fig 7C). In contrast, the 89 late-stage-specific upregulated genes were again predominantly associated with immune functions, specifically type II interferon signaling, negative regulation of innate immune response and innate antiviral immune response (Fig 7D).

Finally, GO analysis of the 60 genes downregulated specifically at the early stage identified enrichment for processes such as multi-organism reproductive process, antigen processing and presentation of peptide antigen via MHC class I, lymphocyte activation, signaling by NTRKs, insulin secretion, thermogenesis, extracellular matrix organization and adrenergic signaling in cardiomyocytes (Fig 7E).

### MAYV detection in reproductive tissues and sperm

Following mating, the liver, spleen, *quadriceps femoris* muscle, ovaries, uterus, seminal vesicles and testes were analyzed for the presence/absence of MAYV by plaque assay in all animals. RT-qPCR was performed on the spleen of infected animals and on the corresponding organs from exposed animals. To detect the presence/absence of MAYV in sperm, RT-qPCR was performed and following cell culture amplification and plaque assay.

Infectious viral particles were detected in the spleen (7/7), ovaries (3/7) and muscle (1/7) of infected females mated with uninfected males. In contrast, all organs analyzed from exposed males were negative by both plaque assay and RT-qPCR. However, after cell culture amplification of sperm samples followed by plaque assay, infectious virus was detected in one exposed male (1/7), which had been paired with the female positive for infectious virus in both the spleen and muscle.

In infected males mated with uninfected females, infectious viral particles were consistently detected in the testes (6/6) and in the spleen (3/6). Plaque assays performed after cell culture amplification of sperm revealed infectious virus in four males (4/6), consistent with the cytopathic effect observed (Fig 8). Viral RNA was also detected by RT-qPCR in direct unamplified sperm samples from the same four animals. No infectious virus was detected in the seminal vesicles. Among exposed females, infectious viral particles tested positive in two animals: the uterus and ovary of one female and the spleen of another, although the RT-qPCR resulted below the limit of detection.

**Figure 8.**
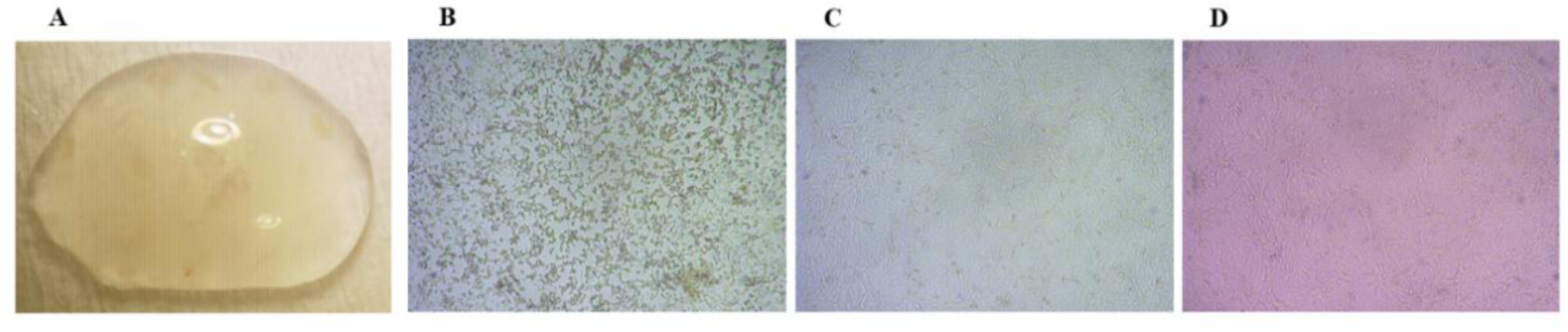
Cytopathic effect of MAYV in sperm cell culture–based amplification. (Representative image). (A) Sperm extracted from the bilateral cauda epididymides. (B‘D) Representative images of Vero cells inoculated with processed sperm samples showing a MAYV-positive sample (B) a MAYV-negative sample (C) and Mock-infected Vero cells (D). Cell morphology was examined by a Zeiss Primovert inverted light microscope (40×).

Overall, viral detection in exposed animals was limited and inconsistent across analytical approaches.

## Discussion

Elucidating the tissue tropism and viral dynamics of MAYV is essential given the limited evidence available for this emerging arbovirus and the relevance of murine models for characterizing infection *in vivo*. Here, we provide evidence of MAYV dissemination to fetal tissues in mice, identifying anatomical targets not previously described. Together, these findings support a model in which gestational timing critically determines placental permissiveness and fetal exposure to MAYV, identifying early gestation as a window of heightened vulnerability during acute infection. Although direct temporal equivalence between murine and human gestation is not possible, early murine gestation (7.5-8.5 dpc) represents a developmental window of active placentation and early organogenesis that broadly parallels the first trimester of human pregnancy, whereas mid-gestation (13.5-14.5 dpc) reflects a more mature placental interface (21,40). Considering that related alphaviruses may cross the placental barrier, our results underscore the importance of including pregnancy models when assessing MAYV transmission routes (41).

To examine early dissemination under controlled conditions, we employed a nanoluciferase-expressing MAYV reporter to evaluate intraperitoneal delivery as a strategy to achieve systemic exposure. *In vivo* imaging showed rapid and sustained viral spread, with peripheral luminescent patterns consistent with the musculoskeletal involvement previously described in experimental models and human disease (18,25,29).

The incomplete overlap between bioluminescent signal and recovery of infectious virus, particularly in older animals, likely reflects differences in analytical sensitivity between reporter-based imaging and plaque assays, the latter requiring a minimum threshold of viable, replication-competent virions, as well as the persistence of viral material following partial clearance. This is consistent with a temporal decrease in viral load over time, suggesting progressive viral clearance despite persistent luminescent signal. In contrast, the consistent recovery of infectious virus in younger mice supports age-dependent variation in susceptibility and viral maintenance within target tissues.

This imaging approach enabled longitudinal, non-invasive monitoring of viral dynamics over a 10-day period, encompassing both acute infection and early recovery phases (24, 25). By providing temporal and anatomical resolution of dissemination events that are difficult to capture in natural transmission settings, the model links viral spread to measurable tissue outcomes.

Together, these observations expand the experimental landscape for MAYV research, supporting mechanistic investigations of tissue tropism, host responses and transmission potential under standardized conditions and offering a framework for future studies addressing reproductive involvement and non-vector routes of spreading.

MAYV WT exhibited rapid and systemic dissemination at 24 and 48 hpi, with infectious virus detected across multiple organs during the acute phase, including detection in reproductive tissues of pregnant females. These results are consistent with other murine alphavirus models, including IFNAR−/− mice, in which widespread viral detection at early time points reflects acute viremia (27). This pattern reflects an initial homogeneous dissemination phase at 24 hpi followed by the emergence of tissue-specific viral accumulation at 48 hpi, consistent with previous murine MAYV models showing that viral burden is influenced by host age, immune status and route of infection, with predominant viral accumulation in skeletal muscle, limbs and lymphoid organs such as the spleen (9, 32). Infectious particles were also detected in muscle, supporting the myopathy and myositis previously described during MAYV infection (9, 24, 28). The increase in viral load in seminal vesicles at 48 hpi further supports progressive involvement of reproductive tissues during acute infection, consistent with the detection of infectious virus in male reproductive organs.

The early hematological shifts observed in our study partially align with previous experimental and clinical reports of MAYV infection. In IFNAR−/− mice model, early leukopenia predominantly driven by lymphopenia was reported during acute infection (28), consistent with the trend toward leukopenia and lymphocyte reduction detected in our model. In contrast, a BALB/c intraplantar infection model described increases in monocytes, neutrophils and lymphocytes during the acute phase (24), indicating that hematological profiles may vary depending on experimental conditions. In human cases, leukopenia, thrombocytopenia and elevated ALT and AST have been documented among reported laboratory abnormalities (17), which parallels the transient cytopenias and aminotransferase elevations observed here. Together, these comparisons suggest that early systemic hematological and biochemical changes are consistent with previously reported features of MAYV infection, although their magnitude and direction may differ across models and clinical settings. Despite the limited mock sample size in pregnant cohorts, the observed shift in leukocyte distribution at 48 hpi across gestational stages consistent with an early systemic inflammatory response rather than a global hematological impairment.

MAYV belongs to the Alphavirus genus (17, 32), which includes viruses capable of affecting the nervous system and embryonic development. The detection of infectious particles in reproductive organs (uterus, ovary, testes, seminal vesicles), with variable detection across tissues at both time points highlights a tissue tropism previously described in immunodeficient mice (14) and raises the possibility of sexual or vertical transmission. Transient and early sex-based differences were observed at 24 hpi. These differences were no longer evident at 48 hpi. While most studies focused on age and nutritional status as determinants of disease outcome (24, 29, 39), these observations suggest a potential sex-dependent component of MAYV infection that warrants further investigation, particularly given similar findings reported for CHIKV (40).

A gestational stage–dependent susceptibility was evident in our model. At the early stage of pregnancy (7.5–8.5 dpc infection; analyzed at 9.5–10.5 dpc), fetuses contained infectious particles at 48 hpi, indicating that MAYV can cross the maternal–fetal barrier during acute infection. In contrast, our data suggest a more restrictive placental environment during mid-gestation (13.5–14.5 dpc; analyzed at 15.5–16.5 dpc), as no infectious particles were recovered from fetal tissues despite significant placental viral loads. It is critical to interpret this’restriction’ specifically as the absence of infectious viral particles detectable by plaque assay. While this confirms a lack of replication - competent virions in the fetus at this stage, it does not preclude the potential presence of viral RNA or non-infectious antigens that might be detectable through more sensitive molecular methods. This is supported by our immunohistochemistry results, where viral antigen was detected in fetal tissues at mid-gestation despite the absence of recoverable infectious virus, suggesting either focal infection below the limit of detection or the persistence of viral antigens following partial viral clearance. This timing-dependent susceptibility is consistent with observations from ZIKV infection, where placental infection and transplacental transmission are strongly associated with maternal infection occurring during the first trimester of human pregnancy, whereas infections acquired at later gestational stages are less frequently linked to fetal exposure (19, 41). This window—analogous to early human embryogenesis (42) — displayed a broad transcriptional response (208 upregulated, 60 downregulated genes), including alterations in developmental and tissue-remodeling programs, as well as downregulation of hormonal signaling, MHC I presentation and ECM organization. These disruptions are compatible with teratogenesis described for alphaviruses such as Semliki Forest virus (39) and ZIKV (20).

In contrast, at mid-gestation infection (13.5–14.5 dpc), infectious viral particles were not detected in fetuses and transcriptional activation was more restricted (139 upregulated, 1 downregulated), dominated by interferon-mediated pathways. This profile aligns with increased placental maturity and antiviral competence (20). The presence of infectious viruses in placenta and uterus, but not in fetal tissues, supports restricted fetal infection at this stage (42, 43). Together, these findings support the concept of a conserved gestational window of vulnerability to vertical transmission among arboviruses, in which early placentation is associated with increased permissiveness at the maternal–fetal interface. The 50 shared upregulated genes across gestational stages—mostly ISGs—reflect a conserved antiviral signature, but timing-dependent transcriptional divergence underscores critical periods of vulnerability, including disruption of angiogenic and developmental pathways (44) and inflammatory responses characteristic of MAYV (24, 25). Viral antigen was detected by immunohistochemistry in placental and fetal tissues at mid-gestation, despite the absence of recoverable infectious virus, suggesting focal or low-level infection below plaque assay detection limits or persistence of viral antigens following partial viral clearance.

While direct extrapolation from murine models to humans is limited, these findings provide a biologically plausible framework to explore potential maternal–fetal risks with MAYV infection in human during pregnancy.

Limitations should be considered when interpreting these findings. Intraperitoneal inoculation does not fully recapitulate natural mosquito-borne transmission and may influence early viral kinetics. However, this route is widely used in murine models of systemic arbovirus infection and was deliberately selected to enable controlled assessment of early systemic dissemination and tissue tropism while minimizing procedure-related distress (45, 46) and local inflammatory confounders. Although intraplantar models better reproduce articular manifestations (24, 25, 27), intraperitoneal delivery provides a standardized framework to evaluate systemic viral dynamics.

Previous murine studies have reported systemic MAYV dissemination and detection of viral RNA in gonadal tissues, suggesting broader tissue distribution (47), although evidence supporting a functional role of reproductive infection in transmission has remained indirect. In the present study, infectious virus was detected in reproductive organs and sperm, supporting the biological plausibility of sexual transmission, though not establishing it as an efficient route and therefore requiring dedicated follow-up studies. Consistent with this, detection among exposed animals was limited and inconsistent, suggesting minimal or transient exposure rather than sustained transmission. If sexual transmission occurs, it is likely to involve low-level infection and restricted replication within gonadal tissues, potentially limiting detectability and contributing to the scarcity of documented cases, a challenge also described for other arboviruses in semen (42).

Taken together, these findings broaden the understanding of MAYV pathogenesis by integrating virological, transcriptomic, histological and imaging approaches within a unified murine model. We provide evidence of expanded tissue tropism, including reproductive and fetal involvement and identify gestational timing as a determinant of placental permissiveness and fetal exposure. Detection of infectious virus in reproductive tissues supports the biological plausibility of non-vector transmission, although its epidemiological relevance remains uncertain. While extrapolation to humans requires caution, these results establish a preclinical framework for investigating MAYV systemic dissemination, maternal–fetal infection and potential transmission routes.

## Conclusion

This study establishes a murine model of MAYV infection that reveals rapid systemic dissemination followed by tissue-specific viral distribution, broad tissue tropism and early involvement of reproductive organs. Transient sex-dependent differences were observed during the early phase of infection, suggesting a potential but limited role of sex in modulating viral dynamics. Gestational stage emerged as a key factor influencing infection outcomes: specifically, infection initiated during early pregnancy (7.5–8.5 dpc) and analyzed at 9.5–10.5 dpc identified a window of increased susceptibility to fetal exposure, associated with the detection of infectious viral loads in fetal tissue. In contrast, infection occurring during mid-gestation (13.5–14.5 dpc) and analyzed at 15.5–16.5 dpc was associated with restricted detection of infectious virus in fetal tissues, despite evidence of viral antigen persistence and sustained viral presence in maternal and placental compartments. The detection of infectious viral particles in reproductive tissues supports the biological plausibility of alternative transmission routes, although their epidemiological relevance remains uncertain. These findings provide a comprehensive experimental framework to advance the understanding of MAYV pathogenesis, systemic dissemination and maternal–fetal involvement and potential non-vector transmission.

## Acknowledgments

We thank José González Santamaría from the Department of Genomics and Proteomics Research, Gorgas Memorial Institute for Health Studies, Panama, for providing the primary antibody used in this study.

## Author contributions

Conceptualization: APA, GM and MC. Methodology: APA, PP, JLP, MPG, BV, JMV, GM and MC. Investigation: APA, PP, JLP, MPG, GG, BV, JMV, GM and MC. Visualization: APA, PP, JLP, MPG, GG, BV, JMV, GM and MC. Statistical Analysis: APA. Supervision: GM and MC. Writing—original draft: APA, PP, JLP, GG, JMV, GM and MC. Writing—review & editing: APA, PP, JLP, MPG, GG, JH, AF, BV, JMV, GM and MC. Project administration: GM and MC. Funding acquisition: GM and MC. All authors agreed to submit the manuscript and have read and approved its final version. Authors take full responsibility for its content.

## Data availability statement

All data supporting the findings of this study are available in accordance with PLOS data sharing policies within the manuscript and its Supporting Information files.

## Supporting information

**S1 Fig. Additional hematological and renal parameters during acute MAYV infection.** Monocytes (A), eosinophils (B), basophils (C), red blood cells (RBC) (D), hemoglobin (E), hematocrit (F), creatinine (G) and blood urea nitrogen (BUN) (H) were measured at 24 and 48 hours post - infection (hpi) and in mock controls. Each dot represents an individual animal; horizontal lines indicate median values. Black symbols denote pre-infection measurements and open symbols denote post-infection measurements. For hematological analyses, n = 12 mice per time point; mock controls, n = 4. For biochemical analyses, n = 6 mice at 24 hpi and n = 4 mice at 48 hpi; mock controls, n = 4. Sample sizes varied depending on the parameter analyzed. Statistical analyses were performed using mixed-effects models (REML) for repeated measures. Differences were considered statistically significant at p < 0.05 (*), p < 0.01 (**) and p < 0.001 (***).

**S1 Table. Differentially expressed genes in placental tissues from MAYV-infected mice at early (7.5–8.5 DPC) and mid (13.5–14.5 DPC) gestation compared with stage-matched controls.**

**S2 Fig. Additional biochemical and metabolic parameters during acute MAYV infection.** Total protein (A), albumin (B), globulin (C), total bilirubin (D), gamma-glutamyl transferase (GGT) (E), blood urea nitrogen/creatinine ratio (BUN/CRE) (F), glucose (G) and body weight (H) were measured at 24 and 48 hours post-infection (hpi) and in mock controls. Each dot represents an individual animal; horizontal lines indicate median values. Black symbols denote pre-infection measurements and open symbols denote post-infection measurements. For hematological analyses, n = 12 mice per time point; mock controls, n = 4. For biochemical analyses, n = 6 mice at 24 hpi and n = 4 mice at 48 hpi; mock controls, n = 4. Sample sizes varied depending on the parameter analyzed. Statistical analyses were performed using mixed-effects models (REML) for repeated measures. Differences were considered statistically significant at p < 0.05 (*), p < 0.01 (**) and p < 0.001 (***).

**S3 Fig. Red blood cell and platelet indices during acute MAYV infection.** Mean corpuscular volume (MCV) (A), mean corpuscular hemoglobin (MCH) (B), mean corpuscular hemoglobin concentration (MCHC) (C), red cell distribution width–coefficient of variation (RDW-CV) (D), red cell distribution width–standard deviation (RDW-SD) (E), mean platelet volume (MPV) (F), platelet distribution width (PDW) (G) and procalcitonin (H) were measured at 24 and 48 hours post-infection (hpi) and in mock controls. Each dot represents an individual animal; horizontal lines indicate median values. Black symbols denote pre-infection measurements and open symbols denote post-infection measurements. For hematological analyses, n = 12 mice per time point; mock controls, n = 4. For biochemical analyses, n = 6 mice at 24 hpi and n = 4 mice at 48 hpi; mock controls, n = 4. Sample sizes varied depending on the parameter analyzed. Statistical analyses were performed using mixed-effects models (REML) for repeated measures. Differences were considered statistically significant at p < 0.05 (*), p < 0.01 (**) and p < 0.001 (***).

**S4 Fig. Exploratory analysis of infection-associated hematological changes across gestational stages.** (A) Neutrophil percentage, (B) lymphocyte percentage, (C) total white blood cell (WBC) count, (D) red blood cell (RBC) count, (E) platelet (PLT) count and (F) procalcitonin (PCT) levels were measured at 48 hpi in females infected at early (7.5 –8.5 dpc) and mid-gestation (13.5–14.5 dpc), compared with their respective mock controls. Data are presented as individual values with median lines. Infected groups: early gestation (n = 6) and mid-gestation (n = 7); mock controls (n = 2 per stage). Statistical analyses were performed using two-way ANOVA with Sidak’s multiple comparisons test. Data distribution was assessed to verify test assumptions. Given the limited sample size of mock controls, statistical comparisons involving these groups should be interpreted with caution. Differences were considered statistically significant at p < 0.05 (*), p < 0.01 (**) and p < 0.001 (***).

